# Integrating physics in deep learning algorithms: A force field as a PyTorch module

**DOI:** 10.1101/2023.01.12.523724

**Authors:** Gabriele Orlando, Luis Serrano, Joost Schymkowitz, Frederic Rousseau

## Abstract

Deep learning algorithms applied to structural biology often struggle to converge to meaningful solutions when limited data is available, since they are required to learn complex physical rules from examples. State-of-the-art force-fields, however, cannot interface with deep learning algorithms due to their implementation. We present MadraX, a forcefield implemented as a differentiable PyTorch module, able to interact with deep learning algorithms in an end-to-end fashion. MadraX is available at madrax.readthedocs.io

---

AlphaFold [1] started a revolution in computational biology by allowing researchers to access reliable structural information for virtually any natural protein rapidly. The general protein folding problem, however, is still far from being solved: while we can predict the conformations of well-studied proteins, inferring the structure of orphan or artificial proteins is still extremely difficult [2]. Similarly, deep learning (DL) algorithms struggle to converge to meaningful solutions in structural tasks if there is not abundant experimental data available [3, 4]: when algorithms have to learn complex physical rules from a limited number of examples, they often result in overfitting. The only currently available solution is to limit the number of trainable parameters in the DL model, thereby reducing the risk of overfitting. Unfortunately, this also limits the learning ability of the algorithm.

An alternative solution to this problem is to define a set of biophysical rules and integrate them directly in the DL algorithm, therefore freeing it from the daunting task of learning physical rules from a few examples. Such sets of biophysical rules are commonly called ‘force-fields’, and they are part of tools such as Rosetta [5], FoldX [6], CHARMM [7] or AMBER [8].

However, traditional force fields are unsuitable for interacting with DL algorithms for two reasons. First, DL algorithms require calculating gradients for the neural network (NN) weights to perform the learning process. This is only possible when the entire DL pipeline is implemented with data-structures that record the gradient of every operation, which is not the case with traditional force-fields. The inclusion of these force fields as such inside a DL algorithm would therefore block the gradient propagation and arrest the training process. In this paper, we refer to the capability of recording the gradient and transferring it to the next step of a DL pipeline as “PyTorch-differentiability” since PyTorch, a very common DL library, provides data structures to handle gradients. Second, DL training is often very computationally intensive; a force-field, therefore, needs to be as efficient as possible both in energy and in gradient calculations to be used in DL settings. This second requirement can be achieved by implementing every physical calculation as operations between tensors, for which functions, highly optimised at the hardware level, are available (i.e. CUDA).

In this paper we present MadraX, a tool which fulfils both requirements. It is a knowledge-based force field implemented as a PyTorch [9] module. We used the FoldX energy function [6], which is a well-established force field for which plenty of validations are available [10, 11], and found a way of transforming its parameters, equations and expressions to make it completely PyTorch-differentiable via the autograd function of PyTorch. Only a few of the terms, for computational and calculation efficiency reasons, were partially modified or integrated (see Methods). MadraX can thus be embedded within any PyTorch NN, like a standard NN layer, maintaining all the properties typical of PyTorch modules.

In the following, we provide a few examples of how MadraX can be used to address common biological problems. We show how MadraX can address very heterogeneous tasks by simply changing the context and its position in the architecture. Additionally, we also show how to combine Madrax with generic optimisation algorithms to solve problems such as protein relaxation and physical simulations.

The purpose of these examples, however, is to show how MadraX can interact with deep learning algorithms at different levels of integration and to illustrate alternative ways of using automatic dif- ferentiation. These examples should, therefore, be considered only as conceptual illustrations of NN architecture implementations, while the development of each of these into mature, adequately trained applications is beyond the scope of this work. Figure 1 shows an overview of the examples we are going to discuss throughout the paper, while Figure 2 shows the computational pipeline in which MadraX has been included to address each of the proposed examples.

**Figure 1:**
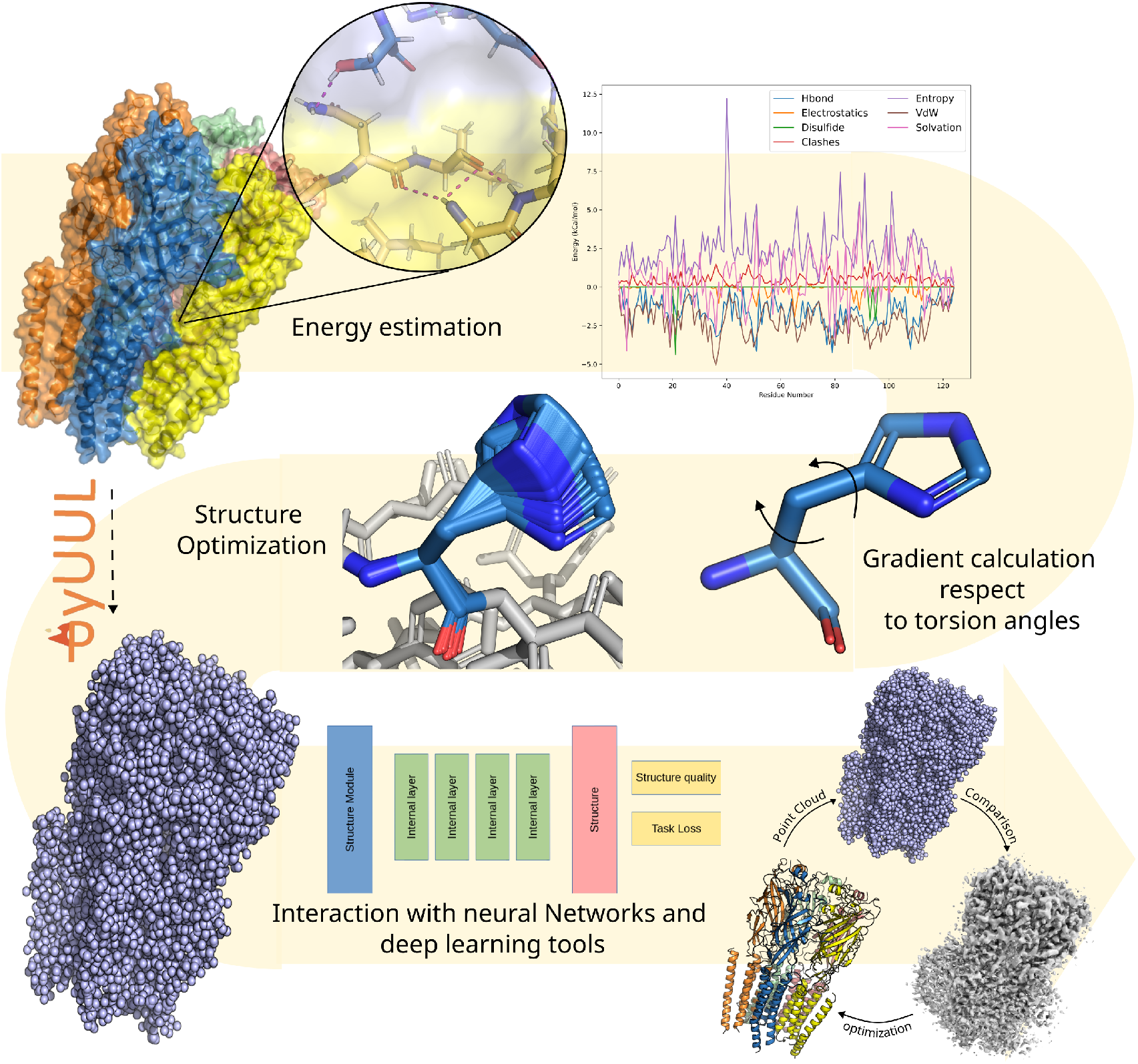
Typical usages of MadraX. MadraX takes a protein structure as input and calculates the energies and atomic interactions that stabilise the fold. It provides the PyTorch-differentiable, tensor-based per-residue energetic landscape of the protein that can directly be used for DL applications. Since MadraX is fully PyTorch-differentiable, it can estimate the conformational energy for protein structure relaxation, calculate the gradient with respect the residues’ torsion angles and apply rotations to find the optimal conformation using gradient descent optimisers. MadraX is implemented as a PyTorch module and can therefore be inserted in any NN and DL tool as easily as any other standard PyTorch NN layer, with the only difference being that MadraX has no trainable parameters. MadraX can be used together with other DL supporting tools, such as PyUUL [12], and it can provide physical constraints to the network learning process, for example, in the refinements of cryo-EM models.

**Figure 2:**
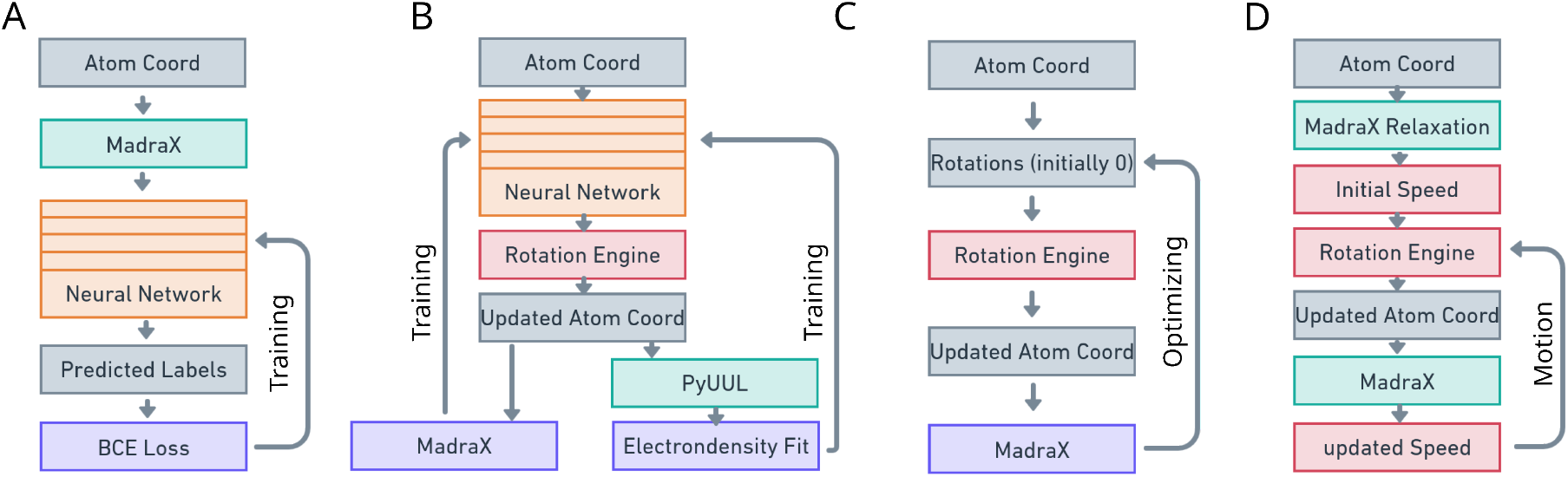
Addressing different problems using different MadraX-based Architectures. MadraX can serve different functions, based on its position in the NN architecture. MadraX was used for mapping small molecule binding sites (A), optimisation of cryo-EM protein structures (B), protein structure relaxation (C), and conformational sampling of protein energy landscapes (D). Orange rectangles represent neural networks, gray rectangles represent simple tensors, red rectangles represent geometrical transformations, green rectangles represent elaboration or data extraction of atomic coordinates, and purple rectangles represent loss functions.

**Figure 3:**
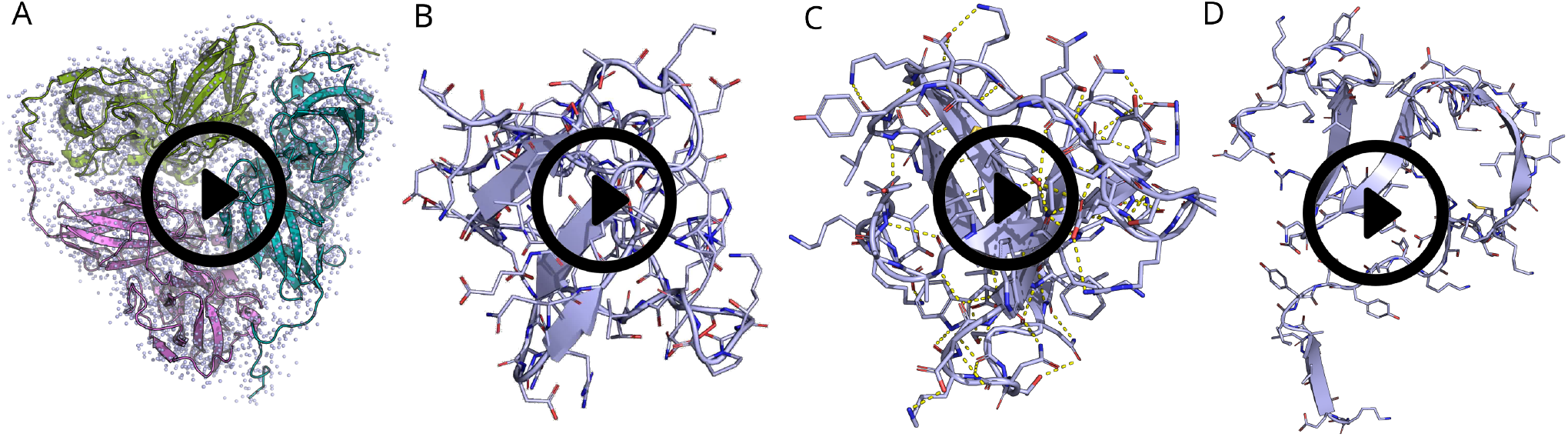
Movies of the three proposed examples. A: The movie shows the optimization of a cryo-EM model in accordance with the similarity between the molecular shape (described by the blue point cloud and calculated with PyUUL) and the experimental electrondensity, coupled with MadraX to maintain physical restraints B: The Movie shows the optimisation of the protein using a PyTorch optimiser coupled with MadraX. The movie shows an optimisation of 50 epochs. C: Pseudo-trajectories calculated with MadraX for human SH3 domain at low temperature. D: Pseudo-trajectories calculated with MadraX for human SH3 domain at high temperature.

## Mapping small molecule binding sites

MadraX can be used to extract the energetical contributions of residues to the conformation of a protein and transform them into a DL-ready tensorial data structure. Since this can be done in just a couple of lines of code, MadraX can be used to build structure-based neural networks with very little effort. One of the possible tasks that can be addressed in this way is the prediction of the residues involved in the binding of small molecules. From [13] we obtained a set of protein structures with annotated binding site residues and then used MadraX to calculate the energetic fingerprint of every residue in the absence of the ligand. We used such energetic fingerprints, along with sequence information, as input of a very simple convolutional NN. Such a network is then trained using a binary cross-entropy (BCE) loss (see Figure 2A for a schematic view of this pipeline). The network converges smoothly, reaching performances comparable to the top-performing methods of predicting residues involved in binding sites (see Supplementary Figure 1). A more detailed discussion of the performances of the network can be found in Supplementary Material section 2.1, while details about the NN are reported in the Methods section. The online MadraX documentation (madrax.readthedocs.io) provides a tutorial about the usage of MadraX in DL algorithms in a non end-to-end fashion.

## Optimisation of cryo-EM protein structures

Since MadraX is PyTorch-differentiable, it can be used in (more complex) end-to-end settings. As a case study, we applied it to the optimisation of protein structures derived using cryo-EM. The experimentally obtained electron-density represents the volume occupied by the atoms of the polypeptide. However, their actual position has to be inferred taking into consideration both the experimental data and the physical constraints. In other words, researchers typically minimise the total energy of the protein structure or complex, to make the model thermodynamically stable, and maximise the match between the resulting total atomic volume and the electron-density.

We used PyUUL, a method we developed earlier to transform protein structures into tensor-based volumetric representations [12], to generate the volume occupied by the atoms of a cryo-EM protein model in a PyTorch-differentiable way. We then built a simple NN which takes the protein atom coordinates as input, it changes/varies the torsion angles (*ϕ, ψ, ω* and *χ* angles) and then generates updated coordinates that are used to calculate volumetric occupancy (calculated with PyUUL) on the one hand and conformational stability (calculated with MadraX) on the other hand. The network is trained to find the torsion angles that produce a protein conformation that has optimal volumetric occupancy of the experimentally determined electron-density map while still respects the biophysical constraints of protein structures. The difference between the volumes of the experimental electron-density map and the suggested protein conformation is compared using Chamfer distance loss (implemented in PyTorch3D [14]), a loss function often used to train NNs in computer vision, while the biophysical component is evaluated by MadraX. Figure 2B shows a schematic overview of the described pipeline.

The final result is a method that iteratively refines the position of the atoms of a cryo-EM model in accordance with the electron-density map in an automated way, without any manual intervention. Movie 3A shows how the network modifies a protein structure and the relative molecular volume during optimisation. A discussion about this application of MadraX is available in Supplementary Information Section 2.3, while details about the NN are reported in the methods section.

## Protein structure relaxation

PyTorch is largely used for the development of NN, and its auto-differentiation algorithms are mostly used to calculate gradients with respect to the NN weights. However, auto-differentiation algorithms can, in principle, calculate derivatives of any arbitrary variable, and this feature can be exploited to perform protein relaxation with MadraX.

More specifically, given the atom coordinates X of a protein P, MadraX can take these coordinates as input and provide an energy E. If we calculate the gradient *δE*/*δX*, we define how E changes depending on the position of the atoms. If we use one of the generic optimisers from PyTorch (developed by e.g. Facebook or Google to address tasks unrelated to biology), we obtain an extremely fast gradient-based structure optimiser to minimise E by moving the coordinates X.

We can also differentiate E with respect to the torsion angles, thereby maintaining the geometry of covalent bonds within the protein. Simply put, this method maintains the FoldX energy function, reformulating the way in which we find the best conformation as a simple and generic gradient-based optimisation problem, a well known and studied computational task for which plenty of algorithms are available. A schematic description of the pipeline is available in Figure 2C. Both these optimisation procedures are purely gradient-based and do therefore not require any protein backbone fragments or side chain rotamer databases as support. Moreover, this application only requires a couple lines of code that is provided as a tutorial in the MadraX documentation.

Movie 3B shows an example of a protein relaxation using this approach after a random perturbation of the position of its atoms. See Methods for more details about the implementation of this procedure and Supplementary Information Section 2.2 for a discussion on the validation of this functionality. MadraX also provides a utility module to implement mutations in the input proteins.

## Conformational sampling of protein energy landscapes

In the previous application regarding protein relaxation, we have been using the automatic gradient calculation in the context of gradient descent optimisers. The numerical calculation of derivatives, however, is also a critical step for the physical simulations. As a final demonstration of the advantage of PyTorch-differentiability, we set up a MadraX-based framework to generate a Molecular Dynamics (MD)-like simulation.

Speed is defined as the derivative of space with respect to time (*δ*X/*δ*t), or, in more practical words, the distance traveled by an object in a small amount of time. In a MD simulation, atoms are assigned with an initial speed and the algorithm simulates the evolution of the system, updating such speed according to the interaction forces between atoms. When calculating the gradient of MadraX’s energy with respect to the atom position, we are estimating how much each atom moves due to the attractive and repulsive forces acting on it in a small amount of time, identified as an optimisation step (or epoch). We can therefore consider it a pseudo speed. Similarly, if we calculate the gradient of MadraX’s energy with respect to the torsion angles, we obtain a pseudo angular speed.

We therefore tried to simulate pseudo motions of atoms by adding, after a relaxation step to ensure the protein is at an energetical minimum, an initial angular speed to each torsion angle in the protein and letting the system evolve by itself. Such initial speed is correlated to the kinetic energy available in the system. We therefore defined a coefficient that we call ‘temperature coefficient’ which describes the average magnitude of the initial motion of atoms. Subsequently, we calculated the gradient of MadraX’s energy with respect to the torsion angles of the protein, obtaining the vector which defines their angular speed. We can simply add this vector to the initial speed to obtain the updated direction and magnitude in which every atom is going to rotate. The position of each atom can be updated as in the previous example using a PyTorch optimiser (in this case we used SGD). We can repeat the same operations, using the new angular speed as initial speed and obtain pseudo-trajectories that evolve in time. The procedure is summarised in Figure 2D.

Movies 3C and 3D show the pseudo-trajectories of a protein using low and high temperature coefficients respectively. In the first one we observe minor local changes, without major conformational shifts, while in the second, one we observe an almost complete protein unfold. This procedure might be used to gather information about the effect of the alteration of the system energy on residue motions, identifying therefore protein regions that are critical for conformational changes.

This example, however, is not meant to replace current state of the art MD methods, since its relationship to physical properties like speed is unclear, The example shows, however, a generalisation of the differentiability concept. As for protein relaxation, this motion simulation does not require any rotamer database: the motion is continuous in space, it is purely gradient-based, and it is possible because MadraX is PyTorch-differentiable.

## Methods

### Definition of the energies

MadraX estimates the energies that stabilise a biological molecule or complex. The energies are defined by hard coding the biophysical formulas in tensorial form and, therefore, no machine learning is involved. MadraX distinguishes/calculates 11 classes of (interaction/stabilisation) energies. Seven of these classes have been taken from the FoldX forcefield [15] and reimplemented in an efficient way in tensorial and PyTorch-differentiable form (see [15] for a detailed description of the Hydrogen Bonds, Electrostatic Interactions, Disulfide Bonds, Van der Waals Energies, Side Chain Entropy and Polar and Hydrophobic Solvation in FoldX). In the next following paragraphs, we will describe how the remaining energies are modelled and calculated.

### Clashes

Clashes occur when two atoms are too close, and their Van der Waals spaces partially overlap. Clashes should almost never be present in high-quality protein models, and atom overpacking is expected to be detrimental to protein stability. MadraX determines clashes and their magnitude based on the minimum distance between specific pairs of atoms observed in high-resolution crystal structures and their atomic radii. To do so, we first downloaded 5000 high-resolution (*<* 1.5*A*) x-ray protein structures from the PDB and for every atom pair within 5A from each other, we collected the difference between their distance and the sum of their radius. We removed from the calculations all the pairs connected by a covalent bond (excluding disulfide bonds) or that were part of a rigid geometrical structure, such as the atoms belonging to the same aromatic ring or the carbonyl and carbon delta of prolines. We also excluded from the calculation atom pairs that were part of the backbone of the same residue. The remaining pairs of atoms were then grouped in six different groups: 1) pairs belonging to the same residue but not directly connected with a covalent bond, 2) hydrogen bonds, 3) disulfide bonds, 4) atoms of the backbone of consecutive residues, 5) donor-acceptor pairs not forming a hydrogen bond, and 6) all the rest. For each group, we calculated the 10th percentile of the distances minus the sum-of-radii difference, which we call correction (see also the formula below). When the clash energy between two atoms is evaluated, we assign an exponentially growing energetic penalty to all the pairs (i,j) associated with a distance that is shorter than the value of its group, using the following formula:

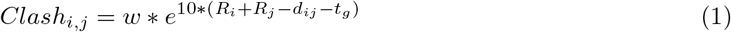

where w is a scaling factor, *R*_*i*_ and *R*_*j*_ are the radii of atoms i and j, respectively, *d*_*ij*_ is the distance between atom i and atom j and *t*_*g*_ is the correction for the group to which the atom pair i, j belongs.

### Backbone Entropy

We calculate the backbone entropy as the log-probability of a residue having specific backbone torsion angles. The distribution of angles was estimated from the high resolution crystal structures already used for the estimation clashes, using a PyTorch-differentiable kernel density estimation (KDE) method based on masked autoregressive flow [16]. For every residue type, we define two distributions: one (with 2 dimensions) for the Phi/Psi angles and one for the Omega angle. The latter has two very narrow Gaussian distributions around -180 and 180 degrees. The final backbone entropy is the sum of these two values. In mathematical terms,

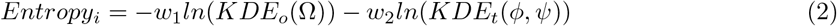

where *w*_1_ and *w*_2_ are two scaling factors, *KDE*_*o*_ and *KDE*_*t*_ are the two mentioned probability distributions and *ϕ,ψ* and *ω* are the backbone torsion angles.

### Peptide Bond Violation

Peptide bonds are covalent interactions between the carbonyl and amino groups of two consecutive residues. Although the geometry of peptide bonds is strongly constrained by atom hybridisation and optimal binding distances, there can be small variations around the optimal values without disrupting protein stability. In order to allow such variability, MadraX gives a penalty to stability based on the deviation from the optimal geometrical values. Specifically, we consider two angles, the CA-N -Cp and CAp-Cp-N, where p indicates an atom of the previous residue, and the distance between the nitrogen and the carbon forming the peptide bond. For each of these values, we built a Gaussian distribution, based on the values observed in the dataset of aforementioned high resolution crystal structures. The final penalty V is then defined as

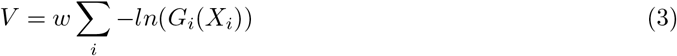

Where w is a scaling factor, i is the estimator under examination (one of the two angles or the bond length), *G*_*i*_ is the Gaussian distribution for the estimator i and *X*_*i*_ is the value of the estimator i for that specific peptide bond.

### Side-chain conformation Violation Penalty

We define the side-chain conformation of a residue by its *χ* torsion angles. Amino acid side-chains have up to 5 *χ* torsion angles. This means that, assuming a fixed covalent bond geometry and length in the side chain, every possible conformation can be defined in a space with up to 5 dimensions, each of which represents a torsion angle. In reality, not all the conformations are physically possible and a large portion of them never or very rarely occur in proteins. To account for this effect, we introduced a side-chain conformation penalty that is calculated using the same KDE approach we used for calculating the backbone entropy, but this time the axis represents the *χ* angles of the amino acid under scrutiny. The probability distribution estimation has been performed using the same dataset of high-resolution crystal structures described in the previous sections. The final penalty for a specific residue is given by the negative logarithmic probability provided by the kernel. In mathematical terms,

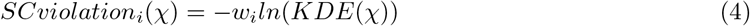

Where *w*_*i*_ is a scaling factor for residue i, KDE is the probability distribution mentioned above for each *chi* torsion angle of the side chain.

### Prediction of residues in a binding site

The network has been trained on the data downloadable from https://zhanggroup.org/COACH/benchmark/. The training set consists of 400 proteins (two of which were removed because of annotation issues), and the testing set contains of 500 proteins. Every residue is annotated with a 1 if it is involved in the binding or a 0 if it is not. The task consists of training a NN on the training set and predicting which residues of the proteins in the test set are involved in binding. For this purpose, we calculated the energetic profile for every protein in the dataset using MadraX, which provides a per-residue estimation of the energies involved as a tensorial representation. We used this tensorial representation, along with the protein sequence, as input for a 3-layer, 1-D convolutional NN with hyperbolic tangent activation, dropout of 0.01 and kernel of 5. The output of this network is taken as input by a 4-layer feed-forward module with 30 hidden neurons, each with hyperbolic tangent activation, dropout of 0.01 and layer normalisation. The network ends in a single neuron, assigning the final probability of interacting with the ligand to each residue. The network has been trained for 200 epochs with Adam optimiser and a learning rate of 0.0001, using a weighted binary cross-entropy with logits as loss function, with weights 1 and 15 for the negative and positive class, respectively, to take into account the class imbalance.

### Cryo-EM model refinement with end-to-end deep learning

The protein complex model and the electron density map file were downloaded from the PDB website (PDB-ID: 6KNF), and the map file was transformed into a Numpy array using the Emda Python package [17]. In this application of MadraX, we have a NN interacting in an end-to-end fashion with the force field module. The network is in charge of generating the set of rotations that minimise the loss functions, and its weights are iteratively refined to reach this goal. The network consists of a feed-forward neural network (FFNN) with 3 layers and Rectified Linear Unit (ReLU) activation. Since we are optimising a single protein and the network is only in charge of “proposing” sets of rotations that minimise the loss functions, it does not take any input. Therefore, the FFNN starts from a single neuron that is always active (always a value of 1). The FFNN ends in a layer containing eight neurons for each residue in the protein, and it defines the set of rotations to perform on each torsion angle of each residue. The backbone is also rotated, and we use the MadraX utility function *madrax*.*mutate*.*rotator*.*RotateStruct* to rotate the starting complex conformation. The newly generated structure is then converted into a volumetric point cloud using PyUUL. Two loss functions are used to evaluate the structure: (1) MadraX itself, that provides the energy of the complex, (2) the Chamfer distance between the PyUUL’s cloud point and the experimental electron-density of the protein complex, turned into a cloud point taking the center of the occupied voxels as points. Occupied voxels have been defined by applying the threshold suggested by the authors of the cryo-EM experiment in the validation report, available on the relative PDB web page.

### Protein relaxation with MadraX

The protein structure used for protein relaxation (Movie 3B) was downloaded from the Protein Data Bank (PDB ID 1AEY). We used the MadraX utility function *utils*.*parsePDB* to parse the file and derive atom coordinates in tensorial form and a list of atom names. We used the atom names to pre-calculate atom information and hashing using the function *dataStructures*.*createinfotensors*. We then created a tensor of shape NxCxRx1×5, where N is the number of protein structures (1 in this case), C is the number of chains and R is the number of residues of the target protein. This tensor represents the rotation applied to the *χ* angles of every residue. The perturbation was implemented as a random rotation (between -0.025pi and 0.025pi for the backbone and -0.25pi and 0.25pi for side chains) applied to every torsion angle of the protein. We estimated the energy of the protein using the MadraX main object Force field, and we calculated the gradient of such energy with respect to the mentioned tensor. We used the PyTorch optimiser object *torch*.*optim*.*Adam* to perform a gradient-descent-based optimisation to find the set of rotations that minimises the protein’s energy, then implemented the rotations using the MadraX utility function *madrax*.*mutate*.*rotator*.*RotateStruct*. The optimisation was run for 100 epochs. This procedure is provided as a tutorial in the online documentation of MadraX

### Motion simulation with MadraX

To simulate the motion of the atoms using a PyTorch setup, we started from the structure of human SH3, downloaded from PDB (PDB ID 1AEY). The positions of the atoms have been parsed using MadraX’s utility function *utils*.*parsePDB*, and we pre-calculated the information required for the energy calculation with the *dataStructures*.*createinfotensors* function. We then relaxed the protein structure, as described in the relevant section above, for 500 epochs with a learning rate of 0.001. The learning rate was reduced during the optimisation using the *ReduceLROnPlateau* PyTorch scheduler with patience of 100 and factor of 0.5. This procedure aims to obtain a conformation as close as possible to a steady state, in which the total energy has a gradient close to 0 with respect to all the torsion angles of the protein. Pytorch, however, uses stochastic numerical optimisers, and this procedure does not completely obliterate the gradient. This means that in our motion simulation, some atoms would start with a higher initial potential energy. To compensate for this, we homogenized the energy states of all residues by adding a starting speed (gradient) so that every torsion angle has same energy. Now that the system is homogeneous from the energetical point of view, we add a Gaussian perturbation to the gradient of every torsion angle. For each torsion angle, the perturbation is sampled from a normal distribution and then multiplied by the so-called temperature coefficient. The temperature coefficient is a scalar that defines the magnitude of the perturbation. We then let the simulation iteratively evolve, and at every iteration we calculate the gradient of the energy calculated by MadraX based on the torsion angles, representing the motion driven by the forces of interaction between the atoms. We add this value to the gradient of the previous iteration, which represents the motion at the beginning of the iteration step. We let the system evolve in this way for 500 epochs. The actual motion is implemented using SGD optimiser from PyTorch, with a learning rate of 10^−7^ and 10^−5^ for backbone and side chain torsion angles, respectively. Movie 3C was generated with a temperature coefficient of 0.015 to simulate low temperature, while for Movie 3D was generated with a temperature coefficient of 15.0 to simulate high temperature.

## Supporting information

supplementary Figures and Sections

