## supplementary Figures and Sections for "Integrating physics in deep learning algorithms: A force field as a PyTorch module"

Orlando et al.

### 1 Supplementary Figures

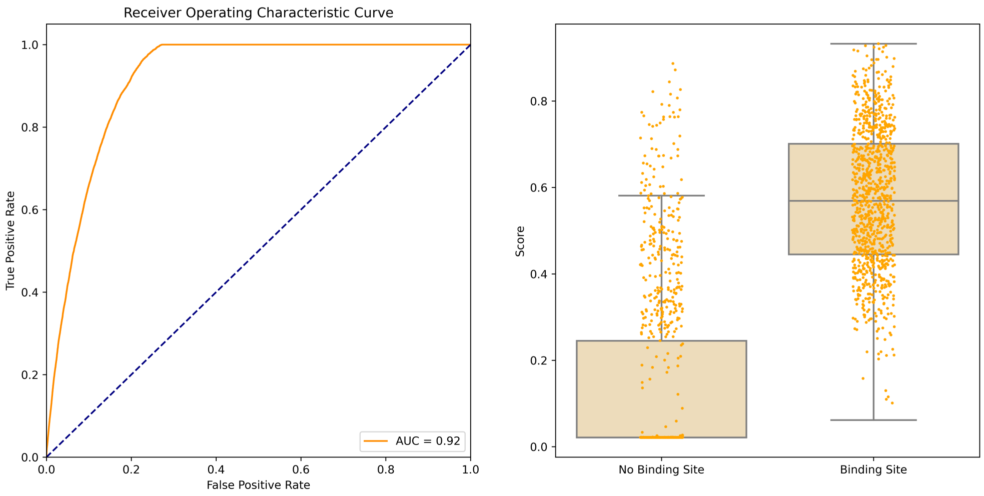


**Supplementary Figure 1**: **Performances of the identification of small molecules binding site**. *Left*, Receiver operating characteristic curve of the presented method on the test dataset. The validation was performed replicating the validation proposed in [1]. The area under the curve (AUC) is 0.92. *Right*, box plots of the scores assigned to residues inside and outside the binding pocket. The orange points show 1000 randomly sampled residues in the two groups and show the actual distribution of the scores.

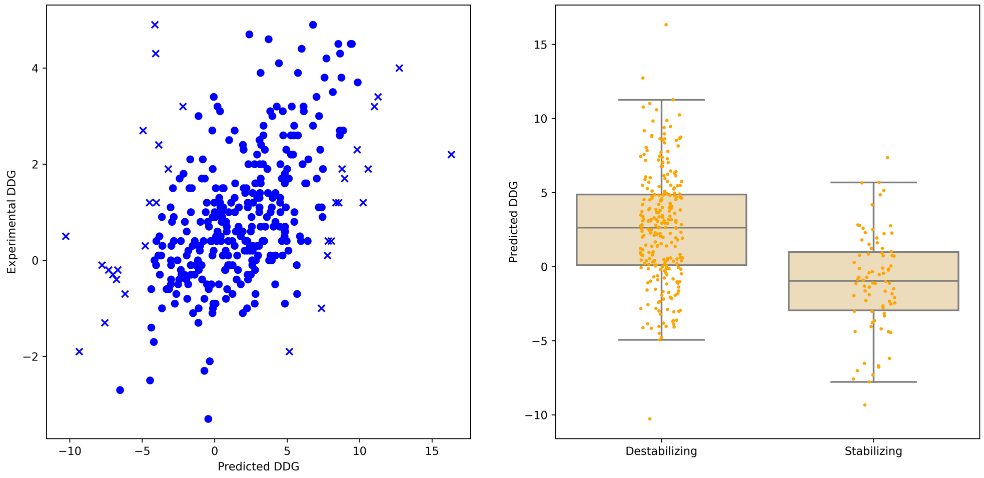


**Supplementary Figure 2**: **Performance of the protein stability prediction task.** *Left*, Plot of the predicted G (horizontal axis) and the experimental $\Delta\Delta$G(vertical axis). The Pearson's correlation coefficient is 0.49. The 10% of the data points in the validation data set that deviates the most from this correlation is marked by x. Right, box plots of the scores assigned to mutations that destabilize and stabilize the protein folding.

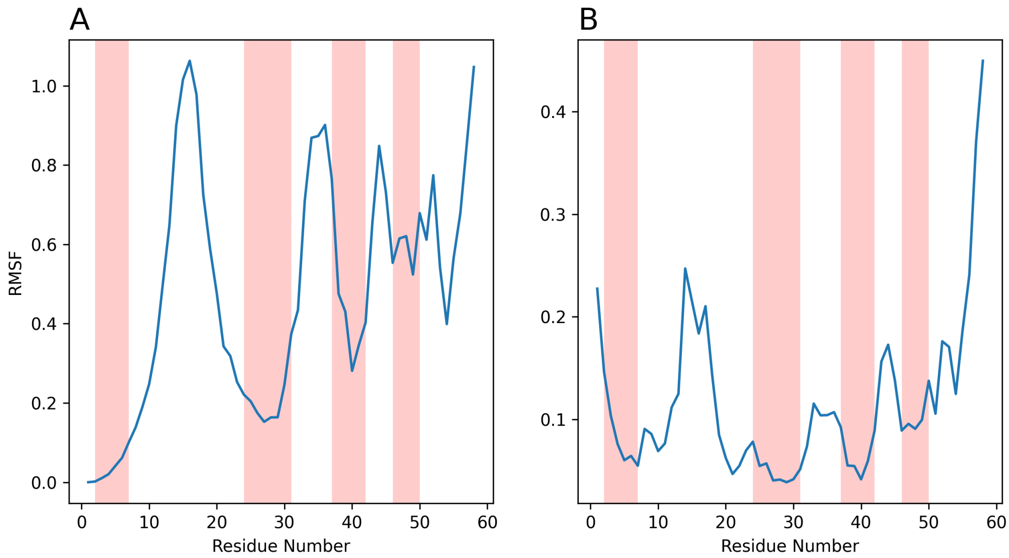


**Supplementary Figure 3**: **The root mean square fluctuation (RMSF) of the simulation of the human SH3 domain**. RMSF of the simulation of the human SH3 domain. The red bars represent the beta strand segments. Panel A shows the RMSF of the unaligned intermediate conformations, while Panel B shows the same results after aligning the intermediate conformations. The figures should be compared with Figure 3 of [1]

### 2 Discussion of performance and validation

#### 2.1 Mapping small molecule binding sites

The prediction of residue-based emergent properties form a typical class of problems in bioinformatics. This is the case with several tasks, such as the prediction of disorder [2, 3], protein aggregation [4, 5] or protein-protein interaction patches [6]. ML methods applied to these tasks often do not use structural data, with a few exceptions [7, 8] that simply extract features such as secondary structure or solvent accessibility from the protein structures. The real complexity of the protein fold, including the forces that stabilize it and the actual interactions between amino acids, is often neglected. To give a straightforward introduction to MadraX, we built a MadraX-based NN that addresses a sequence-oriented task, specifically, the prediction of the residues involved in ligand binding. For validation, we replicated the train-test split approach described in the COACH paper, using the same data and methodology. The final AUC of the validation is 0.92, which means that MadraX outperforms the state-of-the-art predictors reported in the COACH paper [9], ] (the COACH algorithm has an AUC of 0.87).

Supplementary Figure 1 shows the ROC curve and the box-plot of the scores assigned to positively and negatively annotated residues. Given the fact that the datasets are highly unbalanced through the negative class (there are about 15 negatively annotated residues for every positive one), predictors developed in this subject should be considered as prioritization tools instead of classical binary classifiers. Therefore, the goal is to give, on average, a higher score to residues involved in binding, so that the user can focus on a restricted number of residues for experimental analysis. This is why we used a rank-based estimator, such as the AUC, to evaluate the performances.

#### 2.2 Optimisation of cryo-EM protein structures

The goal of this example was to show that MadraX can be used inside a neural network. This would not have been possible with any other force field since, to our knowledge, this is the first pytorch-differentiable force field.

The fact that MadraX is built as a PyTorch module allows us to use it as a loss function for an optimization and combine it with methods typical for computer vision, such as PointNets. This approach shows that it is possible to use machine learning to automate and standardize the process with which atom positions are refined in cryo-EM, taking the raw experimental data into account. During the optimisation of the model, the Chamfer distance gets reduced by 0.128, meaning that the volume of the atomic structure and the electron density map become congruent. At the same time, the Madrax energy was reduced by 25 kCal/mol, showing that the optimization also benefited the stability of the conformation. In this example, we used a very simple neural network architecture since the development of an actual cryo-EM optimization algorithm was outside the scope of this article. To develop such an algorithm, a more sophisticated network architecture would have to be explored. Such an addition might result in a faster and more stable convergence of the network, resulting in a faster and more accurate optimization.

#### 2.3 Protein structure relaxation

To confirm that MadraX maintains the properties and predictive capabilities of a standard force field, we estimated its power to predict the effect of mutations on the stability of proteins. For this purpose, we downloaded a dataset of 342 mutations available on Protherm [10], downloaded from [11], for which a validation using a large number of methods is available [11] (see [11] for the list of mutations taken into consideration), along with the respective wild-type structures.

We implemented the mutations in the respective wild-type structures using MadraX utility function *mutate.mutatingEngine.mutate*. We then defined a NxCxRxMx5 tensor, where N is the number of protein structures, C is the maximum number of chains, R is the maximum number of residues, and M is the maximum number of mutations per protein. This tensor represents the rotation applied to the $\chi$ angles of every residue. We used the MadraX utility function *mutate.rotator.RotateStruct* to implement the rotation. In this function, the mutant and the wild-type protein share all other amino acids with the exception of the mutated ones, and therefore most amino acids are rotated the same way. This gives a huge boost to computational speed because of decreased memory usage and because it reduces the variability in estimation caused by stochastic optimisers. We then used a PyTorch optimizer (Adam) with a learning rate of $10^{-2}$ for 50 epochs to find the set of rotations that minimize the energy of both the mutant- and the wild-type structures to ensure that both are in a conformation associated with an energetic minimum.

To estimate the effect of mutations, we simply took the difference between the energy of mutant and wild-type proteins. The correlation between the predicted $\Delta\Delta$G and the experimental one is 0.49 , which is in line with the bests force field-based stability predictors. Supplementary Figure 2 shows the scatter plot between the predicted and experimental $\Delta\Delta$G. Despite the excellent performance of MadraX this is just an example of the usefulness of MadraX's PyTorch-differentiability, and MadraX should not be considered a predictive tool.

Although the development of a stability prediction tool is outside the scope of this paper, it is technically possible to build a fast and easy-to-use stability predictor with MadraX. Specifically, we think such a tool would primarily benefit from an ad hoc policy to focus on the optimization of the residues that are likely to have a different conformation in the mutant. Such a policy, currently absent in this example, would reduce the number of dimensions the optimizer has to work in, along with the number of differentiation operations that would be required. Protein stability predictors, such as Rosetta or FoldX, never optimize the entire protein because rotating the side chain of every single residue would require such a large amount of rotations that would negatively affect the ability of the optimizer to find the absolute minimum of the energy function. For this reason, limiting the number of rotations the optimizer has to deal with would most probably improve both the speed and the performance of the predictor.

#### 2.4 Conformational sampling of protein energy landscapes

In this example, we showed how to use automatic differentiation to calculate pseudomotions, using the computational properties of MadraX to estimate physical values.

SH3 domain is a relatively small protein, and it has been primarily investigated from the dynamics point of view. See [1] for one of these analyses, in which the authors show the results of MD simulations performed with Gromax. Specifically, they provide the root mean square fluctuation (RMSF) of each residue, which estimates how much every residue moves during a simulation. We performed the same analysis with the pseudo-motions calculated by MadraX (see Supplementary Figure 3 for the results). In our simulation, we assign pseudo angular speed to torsion angles. This means that the first residue is the center of the rotation, and it does not move by definition. Supplementary Figure 3A shows the RMSF of the residues generated with MadraX and Supplementary Figure 3B shows the RMSF obtained by performing a structural alignment of the generated intermediate conformations using Pymol's "fit" function [12]. It is important to note that the structural alignment, while it provides a more realistic estimation of the flexibility of the protein termini, introduces alignment errors and, therefore, adds noise. The plots shown in Supplementary Figure 3 are extremely similar to the ones shown in Figure 3 of [1], demonstrating that there is concordance between our pseudo-motions and the actual molecular dynamics simulation trajectories. We would like to stress the fact this is a proof of concept on how to use automatic differentiation in a physical context, and it should not be interpreted as an alternative to MD methods.
